# *Drosophila* Fog/Cta and T48 pathways have overlapping and distinct contributions to mesoderm invagination

**DOI:** 10.1101/2024.02.02.578601

**Authors:** Uzuki Horo, Nat Clarke, Adam C. Martin

## Abstract

The regulation of the cytoskeleton by multiple pathways, sometimes in parallel, is a common principle of morphogenesis. A classic example of regulation by parallel pathways is *Drosophila* gastrulation, where the inputs from the Folded gastrulation (Fog)/Concertina (Cta) and the T48 pathways induce apical constriction and mesoderm invagination. Whether there are distinct roles for these separate pathways in regulating the complex spatial and temporal patterns of cytoskeletal activity that accompany early embryo development is still poorly understood. We investigated the roles of the Fog/Cta and T48 pathways and found that, by themselves, the Cta and T48 pathways both promote timely mesoderm invagination and apical myosin II accumulation, with Cta being required for timely cell shape change ahead of mitotic cell division. We also identified distinct functions of T48 and Cta in regulating cellularization and the uniformity of the apical myosin II network, respectively. Our results demonstrate that both redundant and distinct functions for the Fog/Cta and T48 pathways exist.

## Introduction

During embryonic development, spatially restricted signaling pathways often operate in tandem to elicit cell behaviors (Manning and Rogers, 2014; Burda *et al*., 2023). Cell shape change, cell motility, and cell division are important cell behaviors that sculpt the embryo (Leptin, 2005; Lecuit *et al*., 2011; Heisenberg and Bellaïche, 2013). These behaviors are often coordinately regulated by multiple signaling inputs that regulate the cytoskeleton. The actomyosin cytoskeleton, composed of filamentous actin (F-actin) and the non-muscle myosin-II motor (myosin-II), is a key force generator that drives cell shape change and motility (Vicente-Manzanares *et al*., 2009). Multiple signaling ligands, receptors, and scaffolds can induce actomyosin activity, which often interface at a key small GTPase, RhoA (Jaffe and Hall, 2005; Buchsbaum, 2007).

*Drosophila* gastrulation has been a powerful system to study the impact of cell signaling on RhoA activity (Rho1 in *Drosophila*). During *Drosophila* mesoderm invagination, the spatially restricted expression of *transcript 48* (*t48*) and *folded gastrulation* (*fog*) are required in parallel for mesoderm invagination (Fig. 1A) (Zusman and Wieschaus, 1985; Costa *et al*., 1994; Kölsch *et al*., 2007; Seher *et al*., 2007). Fog is a GPCR ligand that functions through the G protein-coupled receptors (GPCRs), Mist and Smog, to activate the Gα_12/13_ protein Concertina (Cta) (Parks and Wieschaus, 1991; Costa *et al*., 1994; Dawes-Hoang *et al*., 2005; Manning *et al*., 2013; Kerridge *et al*., 2016; Jha *et al*., 2018). Cta is thought to recruit and/or activate a Rho Guanine nucleotide exchange factor, RhoGEF2, to mediate apical Rho1 activation and cell shape change (Morize *et al*., 1998; Nikolaidou and Barrett, 2004; Kölsch *et al*., 2007; Mason *et al*., 2016). T48 is a transmembrane protein with a PDZ binding motif. T48 is thought to bind RhoGEF2 through RhoGEF2’s PDZ domain (Kölsch *et al*., 2007; Urbansky *et al*., 2016), although how the function of this interaction cooperates with the Cta-RhoGEF2 interaction is poorly understood. The Fog/Cta pathway is required for coordinated, synchronous apical constriction (Parks and Wieschaus, 1991; Sweeton *et al*., 1991; Costa *et al*., 1994; Fox and Peifer, 2007). Furthermore, it is known that Cta is required for timely myosin-II accumulation in cells that initially have a large apical area (Xie *et al*., 2016). In the *Drosophila* germband, Cta is required for medioapical myosin-II assembly downstream of RhoGEF2 (Garcia De Las Bayonas *et al*., 2019). While the Fog/Cta pathway and T48 pathways are known to function in parallel to promote invagination (Kölsch *et al*., 2007), the individual contributions of each pathway to myosin-II accumulation and function are unclear.

**Figure 1.**
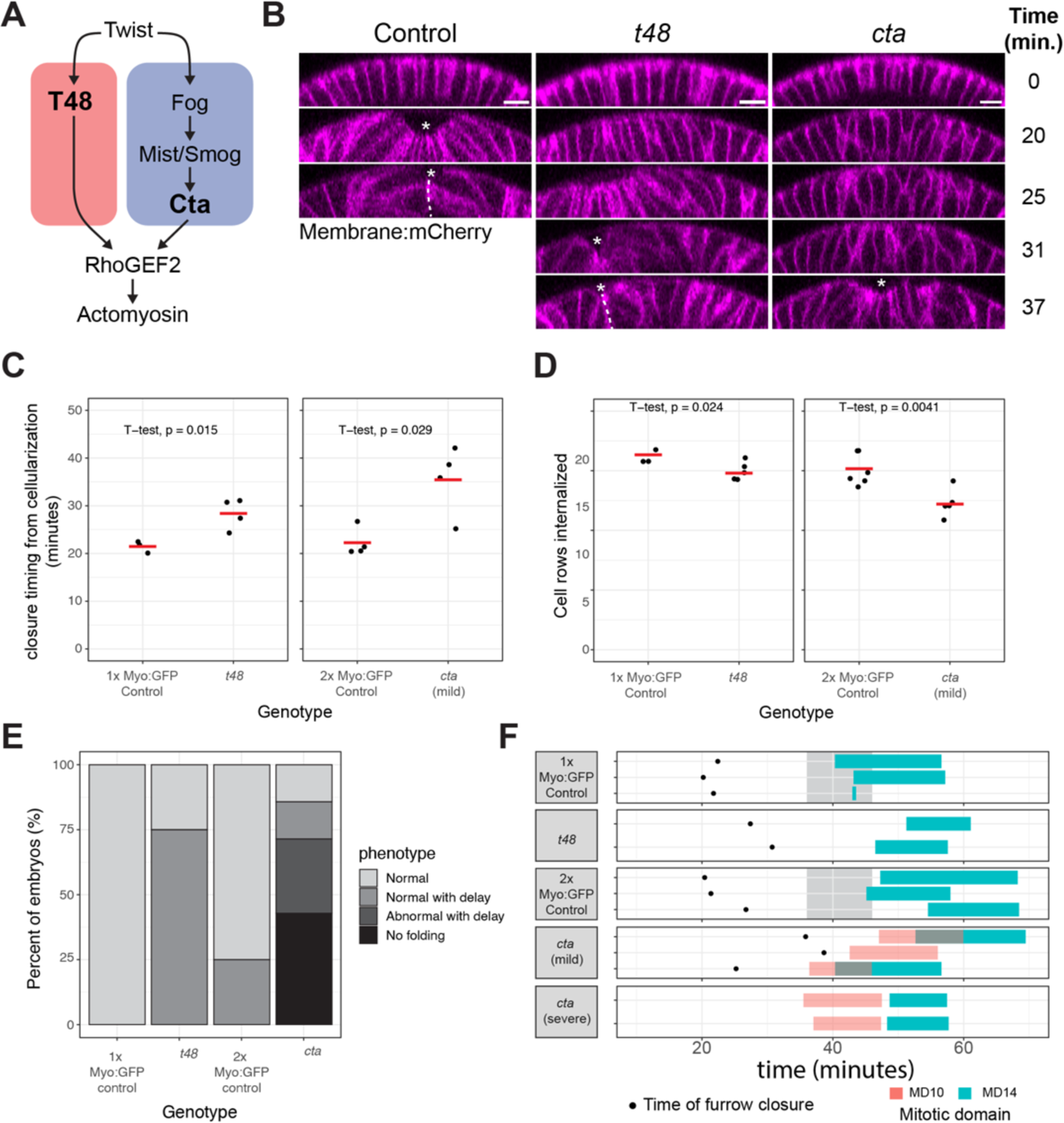
Loss of Cta or T48 pathway delays mesoderm invagination respectively. **(A)** Simplified schematic of the Cta and T48 pathways. **(B)** Mesoderm invagination of control, *cta, t48* mutant embryos expressing Gap43::mCherry (membrane, magenta). Time = 0 at 17 µm cellularization stage. Scale bar, 10 µm. **(C)** Ventral cell rows internalized at the completion of mesoderm invagination in 1x sqh::GFP control (n = 3), sqh::GFP; *t48* (n = 5), 2x sqh::GFP control (n = 6), *cta*; sqh::GFP/sqh::GFP mild embryos (n = 6). *cta* mild embryos had partial mesoderm invagination whereas *cta* severe embryos did not fold. **(D)** The completion of mesoderm invagination relative to 17 µm cellularization stage in 1x sqh::GFP control (n = 3), *t48* (n = 4), 2x sqh::GFP control (n = 4), *cta* mild embryos (n = 4). **(E)** Distribution of mesoderm invagination phenotype in terms of morphology and timing in 2x sqh::GFP control (n = 4), *cta* (n = 7), 1x sqh::GFP control (n = 3), *t48* (n = 4) embryos. **(F)** Timing of the completion of mesoderm invagination, mitotic domain 10 (MD10) divisions, and mitotic domain 14 (MD14) divisions. Time = 0 at 17 µm cellularization stage. In 1x sqh::GFP control, t48, 2x sqh::GFP control, the timing of MD10 division was estimated based on Foe. 1989. Statistical comparison in (C,D) were made with Welch’s unpaired t-test.

Fog and T48 expression are zygotically controlled by the transcription factor, Twist, which is upstream of the transcription factor Dorsal (Costa *et al*., 1994; Kölsch *et al*., 2007). Twist both activates Fog and T48 and delays mesoderm cell division until after invagination via expression of the cell cycle inhibitor, Tribbles (Grosshans and Wieschaus, 2000; Seher and Leptin, 2000). Snail promotes apical myosin II recruitment and cell shape change, in part, by repressing the *Bearded* family of genes (Perez-Mockus *et al*., 2017). Mist expression is regulated by Snail, and modulated by Twist and Dorsal, while Smog is maternally deposited into the embryo (Manning *et al*., 2013; Kerridge *et al*., 2016; Carmon *et al*., 2021). Interestingly, genes downstream of Dorsal and Twist have a dynamic expression pattern – initiating expression at the ventral midline and starting later at more lateral positions (Lim *et al*., 2017; Carmon *et al*., 2021). This ventral-lateral gradient of gene expression is thought to result in a multicellular gradient of T48 protein accumulation, Rho1 activation, and apical myosin II accumulation (Spahn and Reuter, 2013; Heer *et al*., 2017; Lim *et al*., 2017; Denk-Lobnig *et al*., 2021).

Here, we measured the effects of *cta* and *t48* mutants on apical myosin-II dynamics and spatial distribution during mesoderm invagination, in addition to other events in early *Drosophila* embryo development. We found that *cta* and *t48* mutants individually cause a delay in mesoderm invagination relative to mid-cellularization. For *cta*, this delay reflected slower and more discontinuous accumulation of medioapical myosin-II following the end of cellularization. For *t48*, we found that the delay in apical myosin-II accumulation was due, in part, to a delay in cellularization. In severe cases, the myosin-II delay in *cta* mutants resulted in a conflict between apical constriction and cell division timing, which fully disrupted invagination. Overall, we discovered clear, but separate roles for Cta and T48 branches of the Twist pathway, suggesting distinct mechanisms by which each branch of the pathway regulates myosin-II activity while synergizing to contribute to all downstream effects of Twist expression.

## Results

### Cta and T48 pathways both promote timely mesoderm invagination

To determine the function of the Fog/Cta and T48 pathways in mesoderm invagination, we examined the dynamics of tissue furrowing in mutants affecting each pathway. For the Fog/Cta pathway, we used *cta* mutants because *cta* is maternal effect and 100% of embryos laid by *cta* mutant mothers are affected regardless of zygotic genotype (Parks and Wieschaus, 1991). For the T48 pathway, we used *t48* mutants, which are homozygous viable (Kölsch *et al*., 2007), enabling us to collect embryos that are maternal and zygotic *t48* mutant. First, we visualized the timing of invagination relative to cellularization in *t48* and *cta* mutants. We found that both *t48* and *cta* mutants exhibited invagination, but that invagination was delayed (Fig. 1B). We quantified this delay by measuring the closure time – the time when the apical surfaces of cells on adjacent sides of the ventral furrow touched – relative to a benchmark time in cellularization (17 µm furrow canal depth, time = 0). Both *cta* and *t48* mutants significantly delayed invagination compared to matched controls (Fig 1C). Although both mutants caused delays, *cta* mutants had a greater variability and severity in the invagination phenotype - some embryos completely failed to invaginate and others did so, but often with abnormal morphology and fewer cells internalizing (Fig. 1D, E). In contrast, *t48* mutant embryos consistently invaginated, despite the delay (Fig. 1B, D, E). Therefore, *cta* and *t48* mutants both delay mesoderm invagination as single mutants, with the delay in *cta* being most severe.

The *cta* mutants that failed to invaginate (i.e. *cta*, severe) most often exhibited mitotic domain 10 (mesoderm domain) divisions at the embryonic surface, which depleted apical myosin, and reversed apical constriction and tissue furrowing, as observed previously (Fig. 1F, Supplemental Video 1 and 2) (Ko *et al*., 2020). We also observed surface mitotic domain 10 divisions in *cta* mutants with clear invaginations (i.e. *cta*, mild), because the mesoderm failed to fully internalize (Fig. 1F, Supplemental Video 3). The timing of these divisions happened around the time expected for wild-type embryos based on the previous literature (Foe, 1989), at a time when the mesoderm cells would have normally invaginated. We did not assess the timing of domain 10 divisions in W-T and *t48* mutants because our imaging resolution was not sufficient to capture internalized mesoderm tissue, the apical myosin network, and the onset of tissue invagination with sufficient time resolution. Thus, in contrast to *t48* mutants, *cta* mutants sometimes caused a delay that led to a conflict between apical constriction and mitotic domain 10 cell divisions, which disrupted mesoderm internalization, similar to the effects of premature cell division in a cell cycle regulator mutant (Grosshans and Wieschaus, 2000; Seher and Leptin, 2000).

### Cta and T48 pathways are not required for medioapical myosin, but promote timely myosin accumulation

To determine the mechanism that underlies the delay in mesoderm invagination in these mutants, we measured the timing of apical myosin accumulation and its accumulation rate. We found that apical myosin could still accumulate in both *cta* and *t48* mutant embryos (Fig. 2A). Consistent with the invagination delay, both *cta* and *t48* mutants resulted in a slower rate of myosin accumulation, leading to lower total levels of myosin in the mesoderm during invagination (Fig. 2A - C). The defect in myosin accumulation was most severe for *cta*, but the delay was also prominent for *t48*, despite the normal appearance of mesoderm invagination in this mutant. Thus, both maternal *cta* and maternal and zygotic *t48* mutants significantly delayed the rate of myosin accumulation.

**Figure 2.**
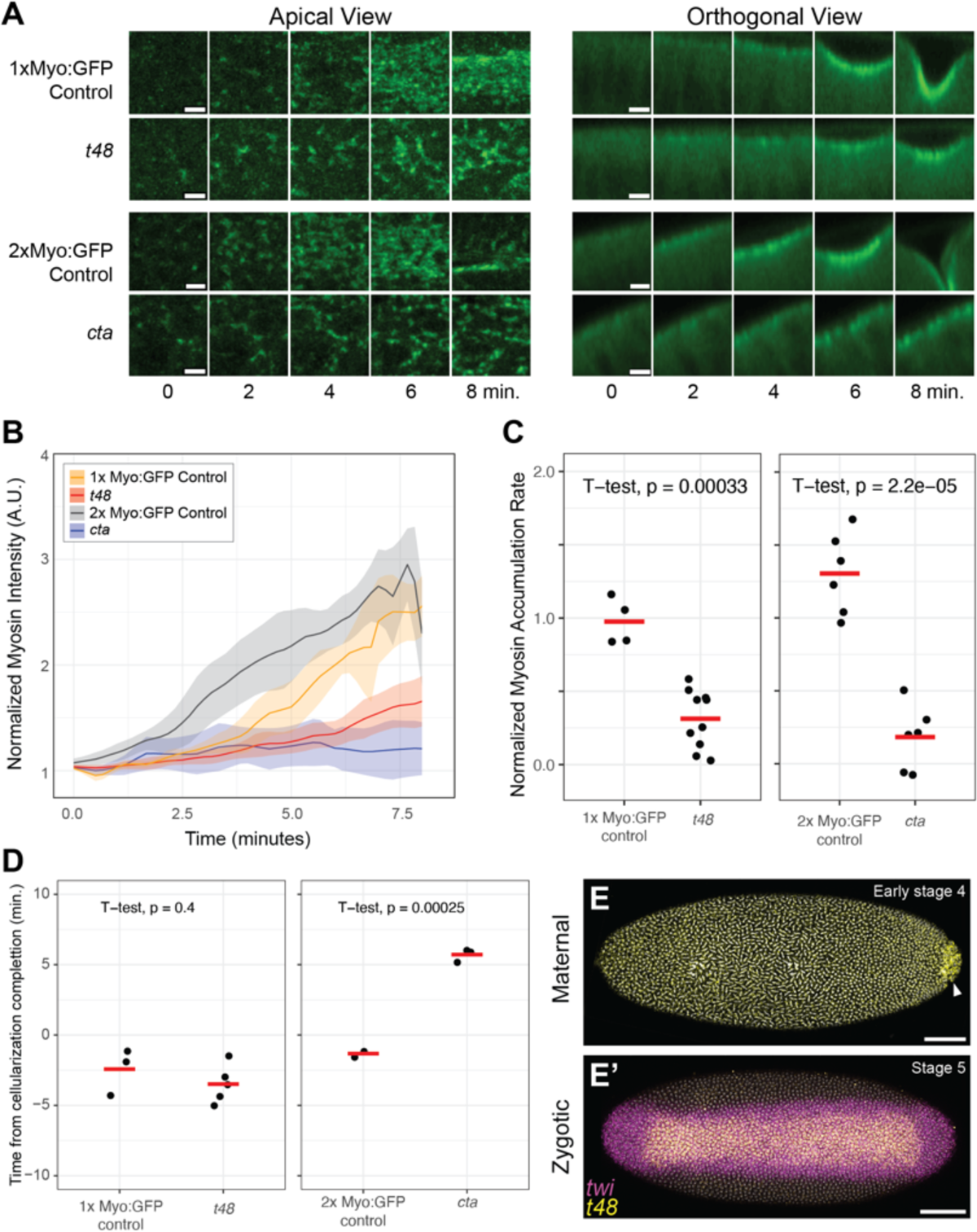
Loss of Cta or T48 pathway reduces the myosin accumulation rate during mesoderm invagination. **(A)** Time-lapse images of myosin accumulation during mesoderm invagination from both apical and orthogonal view. *cta* and its control embryos express two copies of sqh::GFP (Myo:GFP, green), whereas *t48* and its control embryos express one copy of sqh::GFP. Time = 0 at the onset of myosin accumulation. The onset of myosin accumulation was determined as described in the method section. Scale bars: 5 µm. **(B)** Quantification of normalized myosin fluorescent intensity at the ventral midline of two copies sqh::GFP control, *cta* mutant, one copy sqh::GFP control, *t48* mutant embryos. Solid lines represent mean and shaded area represent plus or minus standard deviation. Time = 0 at the onset of myosin accumulation. **(C)** Quantification of normalized myosin accumulation rate during the first 6 minutes of mesoderm invagination in in 1x sqh::GFP control (n = 4) and *t48* (n = 10) embryos (left) and 2x sqh::GFP control (n = 6) and *cta* (n = 5) embryos (right). **(D)** Timing of the onset of myosin accumulation from cellularization completion in 1x sqh::GFP control (n = 3) and *t48* (n = 5) embryos (left) and 2x sqh::GFP control (n = 2) and *cta* (n = 3) embryos (right). Statistical comparison in (C,D) were made with Welch’s unpaired t-test. **(E, E’)** Fluorescent in situ hybridization for twist and t48 mRNA in early stage 4 and stage 5 wild-type embryos. Arrowhead points to pole cells. Scale bar: 50 µm.

By tracking the depth of the cellularization front in confocal movies from control and *t48* mutants, we found that the timing of mesoderm invagination was not delayed relative to a cellularization depth of 30 µm (Fig. 2D). In contrast, *cta* mutants exhibited a clear delay in apical myosin accumulation relative to the end of cellularization (Fig. 2D). Consistent with the earlier role for T48, we found that T48 transcript was ubiquitously present at early stages, as previously described (Urbansky *et al*., 2016). The presence of ubiquitous, possibly maternal, transcript preceded spatially restricted zygotic gene expression in the center of the Twist expression domain (Fig. 2E).

### Myosin accumulation delay in t48 mutants is, in part, a consequence of a cellularization delay

To determine if the delay in apical myosin accumulation reflected a cellularization delay in *t48* mutant embryos, we measured the increase in cell height over time in confocal movies. We found a decreased rate of cellularization in *t48* mutants in comparison to controls (Fig. 3A – B). The cellularization rate in *cta* mutants was more variable than that of control or *t48* mutants, however, on average occurred at similar rate as controls (Fig. 3B). The timing of myosin accumulation onset for *t48* mutants was anti-correlated with the prior rate of cellularization (Fig. 3C), suggesting that the mesoderm invagination delay in the *t48* mutant was due to a delay in cellularization.

**Figure 3.**
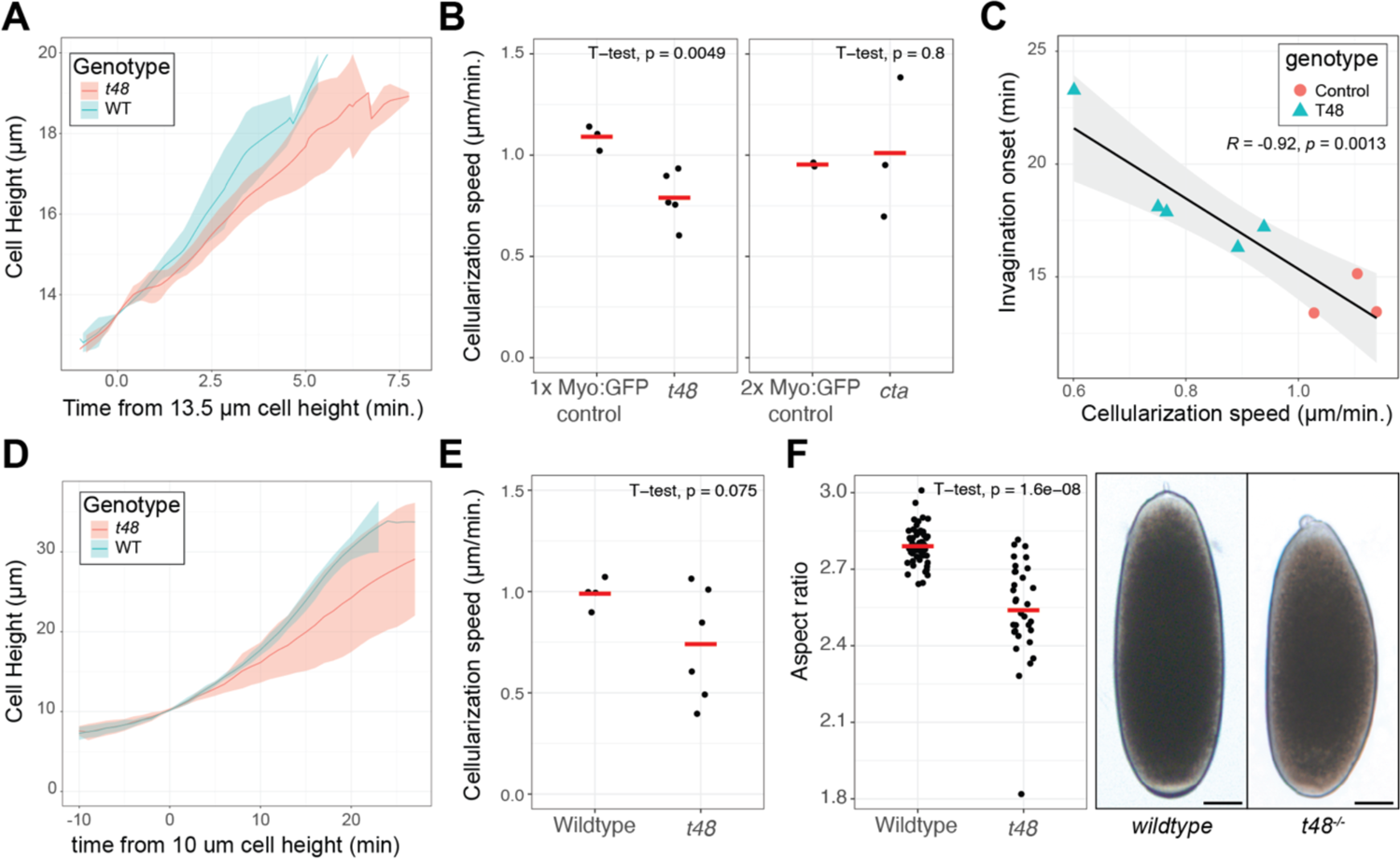
*t48* mutants exhibit the reduced cellularization speed, which correlates with delayed invagination onset. **(A)** Cell height dynamics during cellularization based on sqh::GFP localization at the cellularization front. Solid lines represent mean and shaded area represent plus or minus standard deviation. Time = 0 at 13.5 µm cellularization. **(B)** Quantification of cellularization speed in 1x sqh::GFP control (n = 3) and *t48* (n = 5) embryos. **(C)** Correlation between the cellularization speed and the onset of myosin accumulation from 13.5 µm cellularization in 1x sqh::GFP control (n = 3) and *t48* (n = 5) embryos. R and p are Pearson correlation coefficient and p-value. **(D)** Cell height dynamics during cellularization based on brightfield imaging of cellularization front. Solid lines represent mean and shaded area represent plus or minus standard deviation. Time = 0 at 10 µm cellularization. **(E)** Quantification of cellularization speed in control (n = 4) and *t48* (n = 6) embryos without sqh::GFP. **(F)** Aspect ratio of embryos before or during gastrulation in control (n = 54) and *t48* (n = 33) without sqh::GFP. Images represent brightfield views of early syncytial embryos laid by indicated genotypes. Scale bar: 50 µm. Statistical comparisons in (B,E,F) were made with a Welch’s unpaired t-test.

We verified the cellularization defect in *t48* mutants using stocks that lacked GFP fusion proteins by using brightfield microscopy to visualize furrow canal decent towards the yolk. Cellularization has two phases: an initial slow phase where the furrow descends to ∼ 10 µm depth, and a later fast phase where the furrow canal descends the remaining distance to the yolk (Merrill *et al*., 1988). Tracking the furrow canal position demonstrated that *t48* mutants undergo the initial slow phase normally but failed to speed up during fast phase for an overall slower cellularization rate (Fig. 3D – E). Notably this defect happened around the entire embryo, consistent with the presence of uniformly distributed *t48* transcripts at this stage (Fig. 2E) (Urbansky *et al*., 2016). In addition to the cellularization defect prior to gastrulation, we found that *t48* mutants had rounder egg shape (Fig. 3F). Taken together, these results showed that the T48 pathway has functions prior to gastrulation including regulating cellularization and egg shape.

### Cta and T48 pathways have distinct effects on the spatial distribution of myosin accumulation across the mesoderm tissue

Given the different effects of *cta* and *t48* mutants during cellularization, we next examined whether there were differences between these mutants in the tissue-wide assembly of the apical myosin network. First, we examined the ventral-to-lateral multicellular gradient of apical myosin. By projecting apical-basal cross-sections along the anterior-posterior axis, we could measure the ventral-lateral ‘profile’ of the multicellular apical myosin gradient over time (Fig. 4A – B, Supplemental Video 4). Fitting this myosin intensity profile and measuring the half-maximal myosin width demonstrated that disrupting either *cta* or *t48* reduced the multicellular myosin gradient width (Fig. 4C – D). Furthermore, we measured the fractional apical area coverage by myosin signal, and similarly found that the width of the zone covered by myosin was reduced in both *cta* and *t48* mutants (Fig. 4E – H). Taken together, our data demonstrate that both Cta and T48 are individually required to extend the multicellular myosin gradient laterally into the margins of the mesoderm.

**Figure 4.**
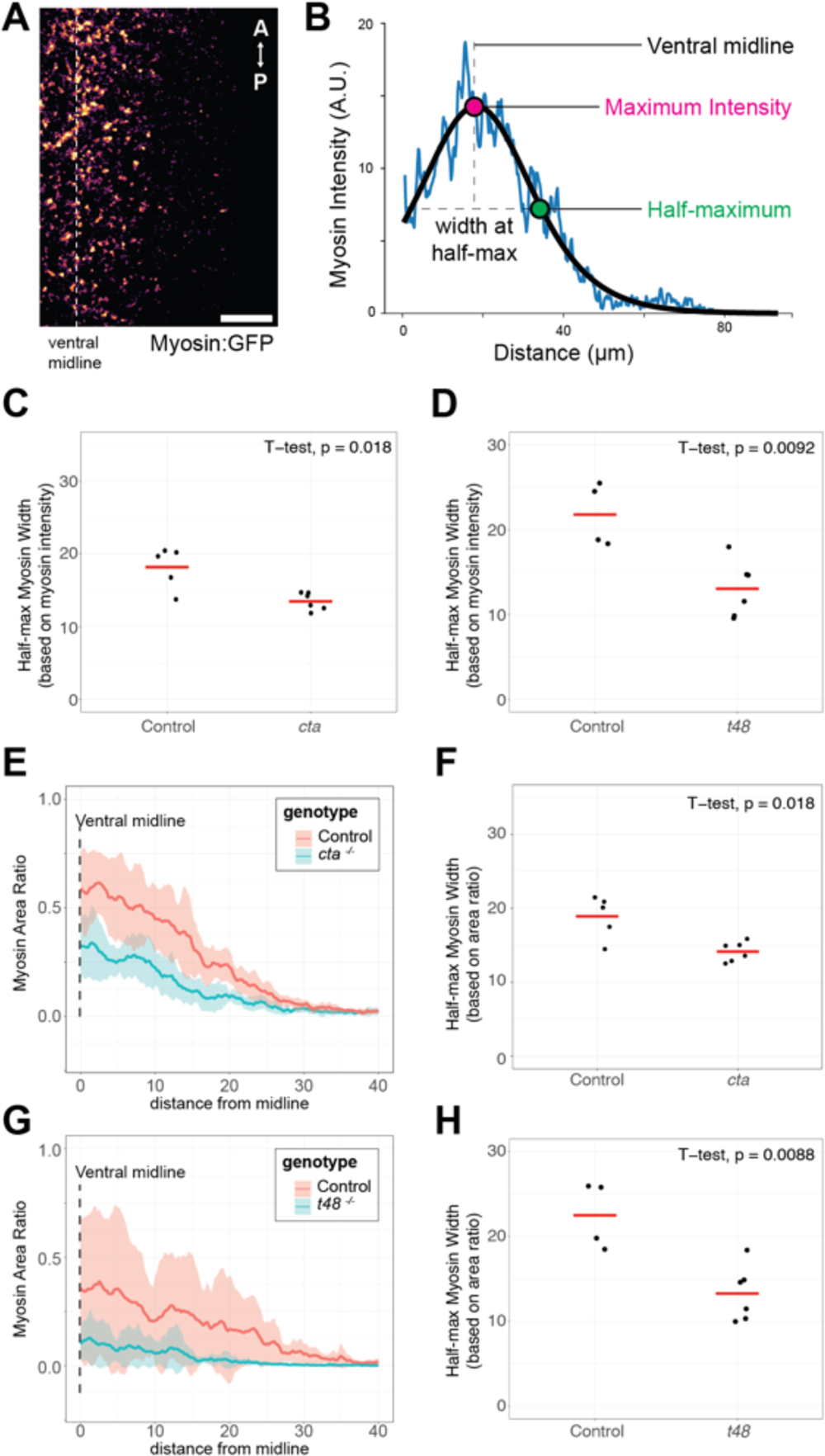
Loss of cta or t48 pathway narrows multicellular mesoderm myosin gradient. **(A)** Representative apical image of myosin accumulation in a 2x sqh::GFP (Myosin:GFP) control embryo at the apical flattening stage. Scale bar: 20 µm. **(B)** Myosin intensity profile from the ventral-lateral axis and a fitted curve of (A). Half-max myosin width is the distance between maximum myosin intensity position and half maximum myosin intensity position of the fitted curve. **(C)** Quantification of the half-max myosin width of 2x sqh::GFP control (n = 5) and *cta* (n = 6) embryos averaged over 2 minutes preceding apical flattening. **(D)** Same quantification as (C) in 1x sqh::GFP control (n = 4) and *t48* (n = 6) embryos. **(E, G)** The ratio of area with myosin accumulation is calculated for the ventral-lateral axis. Solid lines represent mean and shaded area represent plus or minus standard deviation. **(F)** Quantification of the average half-max myosin width based on myosin area ratio in (E). 2x Myo:GFP control (n = 5) and *cta* (n = 6) embryos averaged over 2 minutes preceding apical flattening . **(H)** Same quantification as (F) in 1x Myo:GFP control (n = 4) and *t48* (n = 6) embryos. Statistical comparisons in (C, D, F, H) were made with a Welch’s unpaired t-test.

While we observed that both *cta* and *t48* affected the width of the myosin gradient, we saw a qualitatively different effect of each pathway on the continuity of the supracellular myosin network (Fig. 5A, B, E, and F). To quantify this observation, we measured the porosity of the myosin signal. Both the mean diameter of pores, and the number of large pores were increased in *cta* mutants relative to the control (Fig. 5C – D). Thus, *cta* reduced myosin accumulation while also resulting in a more discontinuous supracellular myosin network. In contrast, *t48* mutants lowered the amount of myosin accumulation without affecting the visible continuity of the supracellular meshwork (Fig. 5 G – H). Taken together, these data showed that while both *cta* and *t48* mutants reduce myosin accumulation rate and the width of the myosin gradient, they affect the organization of the supracellular myosin network differently, likely due to uncoordinated nature of the apical constriction in the *cta* mutant (Sweeton *et al*., 1991; Xie *et al*., 2016).

**Figure 5.**
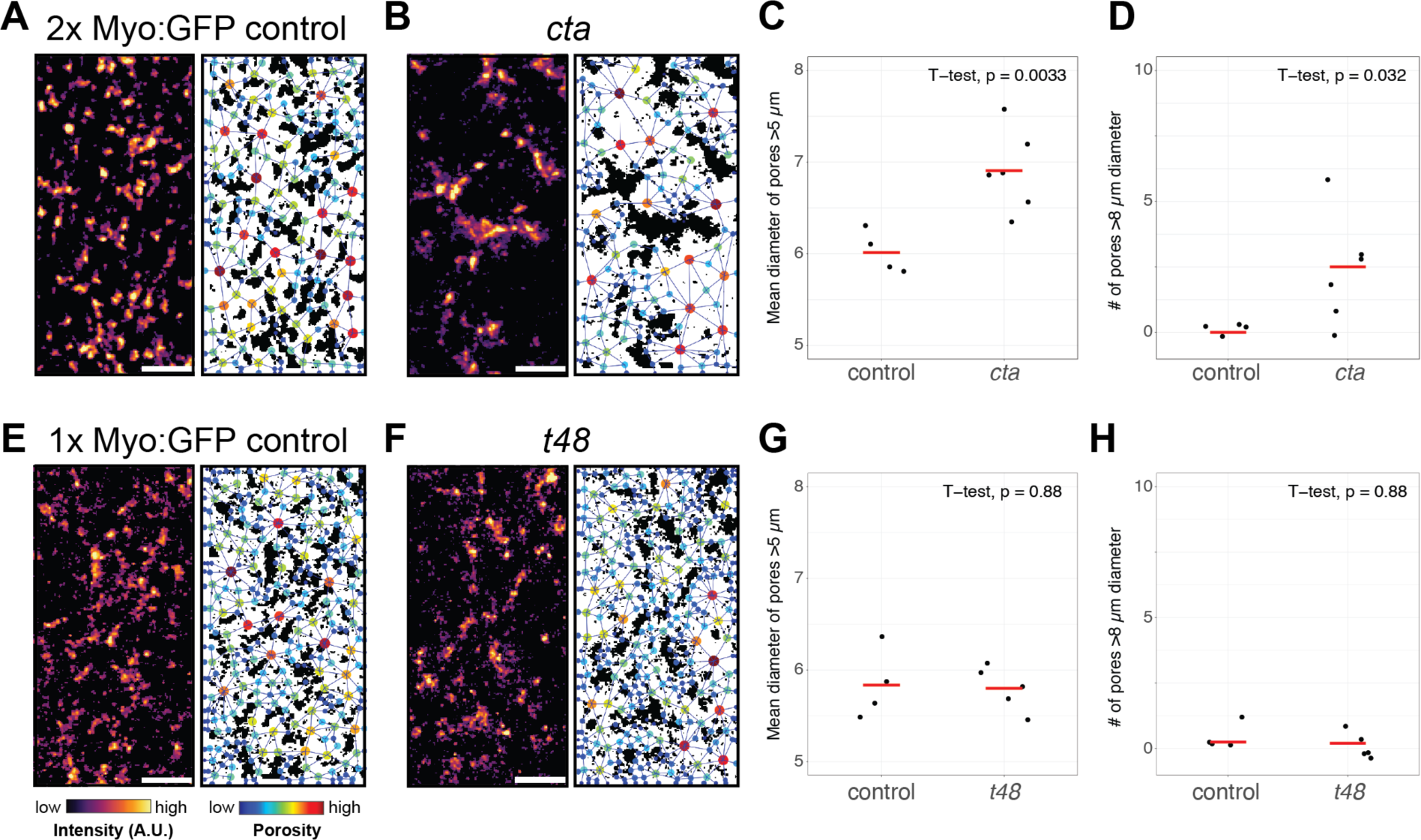
Loss of cta pathway, but not t48 pathway, results in porous supracellular myosin network. **(A, B, E, F)** Representative apical images of supracellular myosin network and pore distribution in 2x sqh::GFP (Myo:GFP) control (A), *cta* (B), 1x sqh::GFP control (E), *t48* embryo (F). Left: myosin distribution at the center of mesoderm (60 um in anterior-posterior axis, 30 µm in dorsal-ventral axis). Right: significant myosin accumulation is present in black regions. Pore networks inferred from white regions are overlayed. The colors of nodes in pore network represent the diameter of each inferred pore. Scale bars: 10 µm. **(C)** The mean diameter of pores greater than 5 µm for each embryo of 2x sqh::GFP control (n = 4) and *cta* (n = 6) at the apical flattening stage. **(D)** Number of pores with greater than 8 µm diameter in each embryo of 2x sqh::GFP control (n = 4) and *cta* (n = 6) embryos at the apical flattening stage. **(G)** Same quantification as (C) for 1x sqh::GFP control (n = 4) and *t48* (n = 5) embryos. **(H)** Same quantification as (D) for 1x sqh::GFP control (n = 4) and *t48* (n = 5) embryos. Statistical comparison in (C, D, G, H) were made with Welch’s unpaired t-test.

## Discussion

The Fog/Cta and T48 pathways function in parallel to promote mesodermal invagination (Kölsch *et al*., 2007). Despite their redundancy in promoting invagination, we found that each pathway had distinct roles in regulating spatial and temporal apical myosin accumulation and cellularization rate. Disrupting the function of either Cta or T48 delayed myosin accumulation and reduced the width of the multicellular myosin gradient, thus neither pathway is sufficient to reproduce the W-T spatiotemporal profile of apical myosin. While both pathways were required for the normal myosin activation, Cta was uniquely required for uniform myosin activation and coordinated apical constriction across the mesoderm (Parks and Wieschaus, 1991; Sweeton *et al*., 1991; Xie *et al*., 2016). Cta was also often required to ensure that cell shape changes and mesoderm invagination preceded mitotic divisions, with *cta* mutants sometimes resulting in a *tribbles*-like reversal of furrowing (Grosshans and Wieschaus, 2000; Seher and Leptin, 2000; Ko *et al*., 2020). In contrast to *cta*, *t48* mutants exhibited uniform, albeit lower levels of, myosin activation and exhibited a reduced rate of cellularization. Therefore, our study demonstrates that the Fog/Cta and T48 pathways are *partially redundant*, with both overlapping and distinct functions during *Drosophila* early embryo development.

### Neither cta, nor t48, mutants resemble twist mutant or fog, t48 double mutant

Twist is required to stabilize apical myosin and maintain constricted cell shape. In the *twist* mutant, cells exhibit apical myosin pulses, but cells fail to transition to a state where myosin accumulates (Xie and Martin, 2015). This failure to stabilize apical myosin is also observed in *fog t48* double knock-down embryos (Martin *et al*., 2010). While *cta* and *t48* mutants delayed myosin accumulation, apical myosin accumulated and promoted apical constriction to the extent that the mesoderm often invaginated. A consequence of failing to stabilize apical myosin is that *twist* mutants fail to assemble a supracellular apical myosin meshwork that spans the mesoderm (Martin *et al*., 2010). The anterior-posterior directed tension that is generated by this meshwork is important for pulling the mesoderm into the embryo (Fierling *et al*., 2022). The *cta* mutant still forms a supracellular apical myosin network, although it is more porous than the normal embryo, presumably because of the uncoordinated apical construction (Parks and Wieschaus, 1991; Xie *et al*., 2016; Yevick *et al*., 2019). The *t48* mutant, despite similarly reducing myosin activation rate, did not result in an uncoordinated apical construction phenotype and assembled a supracellular myosin network. Thus, Fog/Cta and T48 function in parallel to stabilize apical myosin, but have nonredundant functions in promoting timely onset of myosin accumulation. An area of future study will be to understand why the reduced activity through the Twist pathway via Fog/Cta and T48 result in different phenotypes with regards to coordination of apical constriction.

Both *cta* and *t48* mutants reduced the width of the ventral-lateral multicellular myosin gradient, in addition to delaying the onset of accumulation. The timing of *T48* and *mist* transcription onset initiates at nuclei along the ventral midline and progresses laterally (Carmon *et al*., 2021). Furthermore, there is a gradient in the Pol II recruitment rate for each of these transcription units, which leads to graded activity. We observed T48 expression restricted to the central ∼ 12 cells of the mesoderm domain. The expression of Fog has also been shown to exhibit restricted gene expression to the central ∼ 12 cells of the mesoderm, which corresponds to the zone of greatest apical constriction (Costa *et al*., 1994). Our results showed that the combined activity of both T48 and the Fog/Mist/Cta pathways are required for the progression of myosin contractility to marginal mesoderm cells. Any one of these pathways is sufficient for the ventral-most cells to apically constrict, however, both pathways are required for the contractile domain to expand to marginal mesoderm cells.

### The implications of our study with respect to insect evolution

The loss of Fog/Cta pathway or T48 pathway results in the delay of the completion of mesoderm invagination. Interestingly, there seems to be a correlation between the abundance of mesoderm cells and ventral furrow depth in other insects. In *Tribolium*, mesoderm cell distribution and ventral furrow formation is uneven along the anterior-posterior axis. The anterior region has fewer mesodermal cells and forms a shallower ventral furrow than the posterior (Handel *et al*., 2005; Benton *et al*., 2019). In *Chironomus*, *twist*-expressing cells account for around 15% of the embryonic circumference (25% in *Drosophila*) and they form a shallow ventral furrow to internalize mesoderm compared to *Drosophila* (Leptin and Grunewald, 1990; Urbansky *et al*., 2016). The correlation between the width of mesoderm and the depth of ventral furrow during mesoderm internalization among insects suggests a hypothesis that Fog/Cta pathway and T48 pathway were present in the holometabolous insect ancestor, and their activated regions evolved in different insect lineages with different mesoderm distribution before gastrulation. Indeed, *fog* knock-down in *Tribolium* resulted in slower mesoderm internalization and a shallower ventral furrow in the posterior mesoderm (Benton *et al*., 2019). Furthermore, the overexpression of *fog* and *t48* in *Chironomus* increased the number of internalized mesoderm cells (Urbansky *et al*., 2016). Collectively, these studies and our own suggest a conserved function of Fog/Cta and T48 pathways in modulating mesoderm internalization among insects.

In *Drosophila*, the gradients of Fog/Cta and T48 pathway activities from the ventral midline are regulated by the gradient of nuclear Dorsal that contrasts with the uniform expression of *twist* at stages immediately preceding gastrulation (Carmon *et al*., 2021). In *Tribolium*, *fog* expression does not form a gradient from the ventral midline; rather it appears to peak at the periphery of mesoderm (Benton *et al*., 2019). A similar expression pattern is observed in *Chironomus fog2* (Urbansky *et al*., 2016). The peripheral expression pattern of *fog* is counterintuitive because the ventral furrow forms along the ventral midline in both insects. In *Chironomus*, but not *Tribolium, mist* is expressed along the ventral midline which may partly explain how Fog/Mist/Cta pathway activation is restricted to the ventral midline. How *fog* expression is regulated across mesoderm in other insects remain unclear. Although *Toll* knock-down in *Tribolium* results in the loss of *fog* expression in the mesoderm, *twist* knock-down does not change *fog* expression, suggesting a different *fog* expression regulation mechanism from *Drosophila*. Interestingly, the persistence of the Dorsal nuclear gradient during cellularization and early gastrulation is not conserved in *Tribolium* (Chen *et al*., 2000). Dorsal accumulates uniformly across the mesoderm along with weak *twist* expression in the beginning. Then, the Dorsal nuclear gradient gets narrower and ultimately disappears. During ventral furrow formation, only weak cytoplasmic signal is detected in the mesoderm except for the primitive pit. Despite the narrowing of Dorsal nuclear gradient, Twist continues to be expressed across mesoderm during ventral furrow formation (Handel *et al*., 2005). The peripheral expression of *fog* in the mesoderm of basal insects and its relation to the mechanisms mesoderm internalization is an area of future interest.

There are multiple hypotheses on the ancestral function of Fog/Cta pathway and/or T48 pathway. One hypothesis proposes that *fog* and *t48* had ancestral roles in the later epithelial morphogenesis which was then coopted in early development (Urbansky *et al*., 2016). In addition to early development, the Fog/Cta pathway is involved in epithelial morphogenesis in later developmental events such as wing, leg, and salivary gland formation (Nikolaidou and Barrett, 2004; Ratnaparkhi and Zinn, 2007; Manning *et al*., 2013). Furthermore, *fog* and *t48* are present among winged insects whereas their homologs have not been found in non-winged insects to date, implying their ancestral function in later development including wing morphogenesis (Urbansky *et al*., 2016).

Another hypothesis argues that the ancestral function of Fog/Cta pathway is in blastoderm formation (Benton *et al*., 2019). In addition to the abnormal myosin dynamics during mesoderm invagination, we showed that *t48* mutants exhibited slower cellularization speed and rounder egg morphology. Publicly available single-cell transcriptome data shows the expression of *t48* in the somatic cells of the ovariole, suggesting a potential role of T48 in both early embryogenesis and oogenesis (Li *et al*., 2022). Although the involvement of T48 pathway before mesoderm invagination is not known in other insects, Fog/Cta pathway is required for the blastoderm formation in multiple insect species including *Tribolium* (Benton *et al*., 2019). Publicly available *Tribolium* transcriptome data shows the enrichment of *t48* in female gonad albeit in less degree compared to *fog* (Naseem *et al*., 2023). Given that T48 facilitates Fog/Cta pathway during mesoderm invagination in *Tribolium* (Benton *et al*., 2019), it is conceivable that T48 pathway also plays a role in blastoderm formation and/or oogenesis in other basal insects. We showed that in *cta* mutants of *Drosophila*, although the average cellularization speed was not different from control, there is a greater variability. Moreover, previous work showed that *cta* mutants of *Drosophila* show abnormal nuclear position and greater variability in apical cell area before apical constriction (Xie *et al*., 2016). Overall, these studies suggest that the Fog/Cta and T48 pathways are involved in blastoderm formation in *Drosophila* and more basal insects.

## Methods

### Fly stocks and crosses

For *t48* mutants, *t48* was maintained as homozygous stocks with myosin (Sqh::GFP) and membrane (Gap43::mCherry) markers present on the second chromosome. The Sqh::GFP; *t48* flies failed to lay viable embryos. Therefore, embryos from Sqh::GFP/Cyo; *t48* and Gap43::mCherry/Cyo; *t48* flies were used to image myosin and membrane respectively. Single copy myosin controls were embryos from SqhGFP, Gap43::mCherry/Cyo flies.

For *cta* mutants, *cta* was placed over a deficiency (pr31) to examine the maternal effect phenotype of resulting embryos from homozygous *cta* mutant mothers. Briefly *cta*/CyO flies were crossed to pr31/CyO; Sqh::GFP, Gap43::mCherry/TM3 or pr31/CyO; Sqh::GFP. The *cta*/pr31; Sqh::GFP, Gap43::mCherry/Sqh::GFP females or *cta*/pr31; Sqh::GFP/Sqh::GFP flies were collected and crossed to sibling males. Resulting embryos were imaged. For controls with the same copy number of Sqh::GFP, SqhGFP flies were crossed to SqhGFP, Gap43::mCherry/Cyo. SqhGFP/SqhGFP, Gap43::mCherry files were collected and crossed to sibling males. Resulting embryos were imaged.

### Embryonic lethality analysis

Adult flies were allowed to lay eggs for 3-6 hours on apple juice-containing agar plate and the number of eggs were counted. After more than 24 hours, the number of eggs that did not hatch were counted as embryonic lethal. The survival rate of embryo was calculated as 1 – (# of eggs did not hatch / # of total eggs).

### Live imaging (confocal, brightfield)

We collected embryos on apple juice plates and immersed them in Halocarbon 27 oil (Sigma) for staging. We collected blastoderm stage embryos and prepared them for imaging.

For confocal imaging, we dechorionated embryos with 50% bleach, rinsed them with water, and mounted them ventral side up on a slide coated with embryo glue (Scotch tape glue resuspended in heptane, coated onto slide, and allowed to dry). We made a chamber for the glued embryos with No. 1.5 coverslip spacers and a No. 1.0 top coverslip and filled the chamber with Halocarbon 27 oil. We aquired images on a Zeiss LSM710 microscope with an Apochromat 40x/1.2 numerical aperture W Korr M27 objective at room temperature.

For brightfield imaging, we placed embryos with chorion onto a slide with Halocarbon 27 oil and imaged them with brightfield optics using a Nikon 10x/0.25 numerical aperture objective or a Nikon LWD 20x/0.40 numerical aperture objective.

### Image processing and analysis

Images in Figure 1 and 2 were processed using Fiji and those in Figure 4 and 5 were processed using Python 3.9.18. Specific adjustments are described for specific analyses and figure panels below.

### Morphological analysis of mesoderm invagination

In Figure 1 B, membrane channel was processed with a gaussian blur with a radius of 1 pixel and resliced to generate orthogonal views. The timing of 17 µm cellularization, the closure of the furrow and the number of cell rows internalized were manually determined by examining both apical view and orthogonal view images. The morphological phenotype of invagination was manually determined. Embryos which did the closure of the furrow more than 26 minutes later than 17 µm cell height were categorized as delayed (Figure 1 E). Timing of mitotic domain 10 (MD10) and 14 (MD14) divisions were manually determined by examining both apical view and orthogonal view images. Since MD 10 divisions occur in internalized mesoderm in control and *t48* mutant embryos they were not measured and the estimated timing from the literature was used instead. In *cta* mutant embryos, MD14 divisions were distinguished from MD10 divisions based on their relative location in embryo and the division angle. Most MD14 divides in anterior-posterior axis and forms a row of cells, whereas most MD10 divides in apical-basal axis. (Figure 1 F)

### Myosin accumulation during mesoderm invagination

In Figure 2 A, the myosin channel was processed with a Gaussian blur with a radius of 1 pixel. Apical views were obtained by maximum z projection over full stack. To obtain orthogonal views, images were resliced and underwent average Z-projection over 20 µm along anterior-posterior axis. To display myosin intensity differences, contrasts were adjusted by setting the minimum and maximum values. The same adjustments were made for images that were compared.

### Quantification of myosin accumulation at the ventral midline

For analyzing myosin intensity at ventral midline, myosin channel was processed with a gaussian blur with a radius of 1 pixel. Next, images were resliced and underwent average Z-projection over 20 µm along anterior-posterior axis. Then, linear ROIs were drawn to measure myosin intensity profile along apical-basal axis of cells at the ventral midline. The maximum intensity and its position of each myosin profile was obtained. The position was obtained by calculating the position of maximum myosin from that of vitelline membrane. The autofluorescent vitelline membrane position was obtained by finding the first position at which GFP channel intensity is greater than background signal (2.5 was used as a threshold for all embryos).

During mesoderm invagination, myosin intensity is maximum at the apical surface. Therefore, apical myosin intensity was obtained from the maximum myosin intensity. For each embryo, three apical myosin intensity were obtained from different ROIs and averaged to obtain final apical myosin intensity.

In Figure 2, images were aligned at the onset of apical myosin accumulation during mesoderm invagination. The onset of myosin accumulation was defined as the timing when myosin intensity at the apical surface became greater than mean cytoplasmic myosin intensity plus two standard deviation. Mean cytoplasmic myosin intensity and standard deviation was obtained from a rectangle ROI selecting cytoplasm for each embryo. The apical myosin intensity was then normalized to the threshold for quantification (Figure 2 B - D).

### Cell height measurements during cellularization

For cellularization analysis using confocal microscopy, embryos with less than 14 µm cell height at the start of image acquisition and exhibited at least 5 µm increase in cell height were used to capture enough cellularization dynamics. The position of maximum myosin intensity during cellularization was used as the cell height of embryos. This is because during cellularization, myosin intensity is maximum at the cellularization front. (Figure 3 A, B)

For cellularization analysis using brightfield microscopy, embryos with less than 10 µm cell height at the start of image acquisition were used to capture cellularization dynamics. The cell height of the initial frame was manually measured and the movement of cellularization front was tracked by Manual Tracking plugin in Fiji. Measurement and tracking were done three times per embryo and the average cell height at each frame was calculated using R script (Figure 3 D).

### Aspect ratio of embryos

To measure aspect ratio of embryos, brightfield images were converted to 8-bit. To segment embryos, 0 to 200-pixel value was used as threshold. Aspect ratio of segmented embryos was measured using “Analyze Particles” in Fiji with the option “size=100-Infinity” to exclude debris (Figure 3 F).

### Myosin network extraction

Custom Python and R scripts were used to analyze myosin gradient width and pore characteristics of myosin network (Figure 4). Images were first rotated to have ventral side en face and processed with a Gaussian filter with 1 pixel size. To extract myosin network, cytoplasmic background signal was subtracted by setting pixel values with smaller than mean plus two standard deviation cytoplasmic myosin intensity to zero. To obtain cytoplasmic myosin intensity, image frame with no myosin accumulation was manually selected, processed with a gaussian filter with 3 pixel size and background signal (smaller than 2.5 pixels) was subtracted. Two-dimensional myosin network was obtained by conducting maximum intensity Z-projection of the extracted myosin network (Figure 4A). The myosin networks at the apical flattening stage were used in Figure 4 and 5.

### Myosin intensity profile

Myosin intensity profiles along the ventral-lateral axis were generated by averaging myosin intensity of two-dimensional myosin network over anterior-posterior axis (Figure 4B). To quantitatively compare the gradient of myosin profile, the myosin profiles were fit to a function as in (Denk-Lobnig *et al*. 2021): 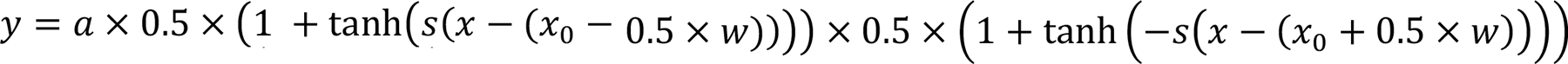, where y corresponds to myosin profile and x corresponds to the position in ventral-lateral axis. Optimal variables were found using curve fit function of scipy.optimize in scipy.1.11.2. Upper and lower limits and initial guesses for each variable were as follows: a: 0, 255, 10; s: 0, 30, 3; w: 0, infinity, 20; x0: -infinity, infinity, manually specified value compatible with the position of ventral midline of each embryo. The distance between maximal myosin intensity and half-maximal myosin intensity of the fitted function was used as the width of myosin gradient. To smooth the fluctuations in myosin gradient width over the time course, the half-maximal myosin width at a given time was averaged with the time steps before and after.

### Myosin area ratio profile

Myosin area ratio profile from ventral midline to the lateral was obtained by calculating the ratio of pixels with myosin accumulation over anterior-posterior axis (Figure 4E, G). Myosin gradient widths were found by fitting the same function as used for myosin intensity profile with the following initial guesses for each variable: a: 1, s: 3, w: 20, x0: the same manually specified value as in myosin intensity profile. To smooth the fluctuations in the myosin gradient over the time course, the half-maximal myosin width of at a given time was averaged with the time steps before and after.

### Myosin network pore size and distribution

To measure size and distribution of pores in myosin network, we used PoreSpy 2.3.0 package (Gostick *et al*. 2019) in Python 3.9.18. Myosin network images that cover at least 15 µm from ventral midline to both left and right sides were selected. To remove the effect of myosin covered area ratio on pore size, myosin network images were binarized with a new threshold such that myosin covered area ratio equaled to 0.25 at the apical flattening stage of each embryo. To detect pores in binarized myosin network images, the SNOW algorithm implemented in PoreSpy was applied with voxel size = 1. (https://porespy.org/examples/networks/tutorials/snow_basic.html). The identified pore network was overlayed on the myosin network images using visualization functions of PoreSpy.

### Statistics

To statistically compare means of two independent groups whose variance were not assumed to be equal, Welch two sample t-test were performed using ggplot2-3.4.4. Normality of sample distribution was tested using Shapiro-test in R.

### Fluorescent *in situ* hybridization

To detect *twist* and *t48* RNA transcripts, we used the hybridization chain reaction (HCR) method, version 3.0 (Choi *et al*., 2018). We designed ∼30 probe pairs for each gene targeting the coding sequence and portions of the UTRs, depending on gene length. Probes were ordered as pooled libraries (OligoPools, Integrated DNA Technologies) and resuspended in nuclease-free water at a concentration of 0.5 µM. Alexa-Fluor-conjugated detection hairpins were acquired from Molecular Instruments. HCR in situ hybridization was performed according to the manufacturer’s protocol using standard formaldehyde-methanol fixed embryos. Following staining, embryos were post-fixed for 15 minutes in 4% paraformaldehyde, and mounted in AquaPolymount. Expression patterns were validated against previously published patterns for both genes.

## References

1. Benton, MA, Frey, N, Nunes da Fonseca, R, von Levetzow, C, Stappert, D, Hakeemi, MS, Conrads, KH, Pechmann, M, Panfilio, KA, Lynch, JA, et al. (2019). Fog signaling has diverse roles in epithelial morphogenesis in insects. Elife 8, e47346.

2. Buchsbaum, RJ (2007). Rho activation at a glance. Journal of Cell Science 120, 1149– 1152.

3. Burda, I, Martin, AC, Roeder, AHK, and Collins, MA (2023). The dynamics and biophysics of shape formation: Common themes in plant and animal morphogenesis. Dev Cell 58, 2850–2866.

4. Carmon, S, Jonas, F, Barkai, N, Schejter, ED, and Shilo, B-Z (2021). Generation and timing of graded responses to morphogen gradients. Development 148, dev199991.

5. Chen, G, Handel, K, and Roth, S (2000). The maternal NF-kappaB/dorsal gradient of Tribolium castaneum: dynamics of early dorsoventral patterning in a short-germ beetle. Development 127, 5145–5156.

6. Choi, HMT, Schwarzkopf, M, Fornace, ME, Acharya, A, Artavanis, G, Stegmaier, J, Cunha, A, and Pierce, NA (2018). Third-generation in situ hybridization chain reaction: multiplexed, quantitative, sensitive, versatile, robust. Development 145, dev165753.

7. Costa, M, Wilson, ET, and Wieschaus, E (1994). A putative cell signal encoded by the folded gastrulation gene coordinates cell shape changes during Drosophila gastrulation. Cell 76, 1075–1089.

8. Dawes-Hoang, RE, Parmar, KM, Christiansen, AE, Phelps, CB, Brand, AH, and Wieschaus, EF (2005). folded gastrulation, cell shape change and the control of myosin localization. Development 132, 4165–4178.

9. Denk-Lobnig, M, Totz, JF, Heer, NC, Dunkel, J, and Martin, AC (2021). Combinatorial patterns of graded RhoA activation and uniform F-actin depletion promote tissue curvature. Development 148, dev199232.

10. Fierling, J, John, A, Delorme, B, Torzynski, A, Blanchard, GB, Lye, CM, Popkova, A, Malandain, G, Sanson, B, Étienne, J, et al. (2022). Embryo-scale epithelial buckling forms a propagating furrow that initiates gastrulation. Nat Commun 13, 3348.

11. Foe, VE (1989). Mitotic domains reveal early commitment of cells in Drosophila embryos. Development 107, 1–22.

12. Fox, DT, and Peifer, M (2007). Abelson kinase (Abl) and RhoGEF2 regulate actin organization during cell constriction in Drosophila. Development 134, 567–578.

13. Garcia De Las Bayonas, A, Philippe, J-M, Lellouch, AC, and Lecuit, T (2019). Distinct RhoGEFs Activate Apical and Junctional Contractility under Control of G Proteins during Epithelial Morphogenesis. Curr Biol 29, 3370–3385.e7.

14. Grosshans, J, and Wieschaus, E (2000). A genetic link between morphogenesis and cell division during formation of the ventral furrow in Drosophila. Cell 101, 523–531.

15. Handel, K, Basal, A, Fan, X, and Roth, S (2005). Tribolium castaneum twist: gastrulation and mesoderm formation in a short-germ beetle. Dev Genes Evol 215, 13–31.

16. Heer, NC, Miller, PW, Chanet, S, Stoop, N, Dunkel, J, and Martin, AC (2017). Actomyosin-based tissue folding requires a multicellular myosin gradient. Development 144, 1876–1886.

17. Heisenberg, C-P, and Bellaïche, Y (2013). Forces in tissue morphogenesis and patterning. Cell 153, 948–962.

18. Jaffe, AB, and Hall, A (2005). Rho GTPases: biochemistry and biology. Annu Rev Cell Dev Biol 21, 247–269.

19. Jha, A, van Zanten, TS, Philippe, J-M, Mayor, S, and Lecuit, T (2018). Quantitative Control of GPCR Organization and Signaling by Endocytosis in Epithelial Morphogenesis. Curr Biol 28, 1570–1584.e6.

20. Kerridge, S, Munjal, A, Philippe, J-M, Jha, A, de las Bayonas, AG, Saurin, AJ, and Lecuit, T (2016). Modular activation of Rho1 by GPCR signalling imparts polarized myosin II activation during morphogenesis. Nat Cell Biol 18, 261–270.

21. Ko, CS, Kalakuntla, P, and Martin, AC (2020). Apical Constriction Reversal upon Mitotic Entry Underlies Different Morphogenetic Outcomes of Cell Division. Mol Biol Cell 31, 1663–1674.

22. Kölsch, V, Seher, T, Fernandez-Ballester, GJ, Serrano, L, and Leptin, M (2007). Control of Drosophila gastrulation by apical localization of adherens junctions and RhoGEF2. Science 315, 384–386.

23. Lecuit, T, Lenne, P-F, and Munro, E (2011). Force generation, transmission, and integration during cell and tissue morphogenesis. Annu Rev Cell Dev Biol 27, 157–184.

24. Leptin, M (2005). Gastrulation movements: the logic and the nuts and bolts. Dev Cell 8, 305–320.

25. Leptin, M, and Grunewald, B (1990). Cell shape changes during gastrulation in Drosophila. Development 110, 73–84.

26. Li, H, Janssens, J, De Waegeneer, M, Kolluru, SS, Davie, K, Gardeux, V, Saelens, W, David, FPA, Brbić, M, Spanier, K, et al. (2022). Fly Cell Atlas: A single-nucleus transcriptomic atlas of the adult fruit fly. Science 375, eabk2432.

27. Lim, B, Levine, M, and Yamazaki, Y (2017). Transcriptional Pre-patterning of Drosophila Gastrulation. Curr Biol 27, 286–290.

28. Manning, AJ, Peters, KA, Peifer, M, and Rogers, SL (2013). Regulation of epithelial morphogenesis by the G protein-coupled receptor mist and its ligand fog. Sci Signal 6, ra98.

29. Manning, AJ, and Rogers, SL (2014). The Fog signaling pathway: insights into signaling in morphogenesis. Dev Biol 394, 6–14.

30. Martin, AC, Gelbart, M, Fernandez-Gonzalez, R, Kaschube, M, and Wieschaus, EF (2010). Integration of contractile forces during tissue invagination. J Cell Biol 188, 735– 749.

31. Mason, FM, Xie, S, Vasquez, CG, Tworoger, M, and Martin, AC (2016). RhoA GTPase inhibition organizes contraction during epithelial morphogenesis. J Cell Biol 214, 603– 617.

32. Merrill, PT, Sweeton, D, and Wieschaus, E (1988). Requirements for autosomal gene activity during precellular stages of Drosophila melanogaster. Development 104, 495– 509.

33. Morize, P, Christiansen, AE, Costa, M, Parks, S, and Wieschaus, E (1998). Hyperactivation of the folded gastrulation pathway induces specific cell shape changes. Development 125, 589–597.

34. Naseem, MT, Beaven, R, Koyama, T, Naz, S, Su, S-Y, Leader, DP, A Klaerke, D, Calloe, K, Denholm, B, and Halberg, KV (2023). NHA1 is a cation/proton antiporter essential for the water-conserving functions of the rectal complex in Tribolium castaneum. Proc Natl Acad Sci U S A 120, e2217084120.

35. Nikolaidou, KK, and Barrett, K (2004). A Rho GTPase signaling pathway is used reiteratively in epithelial folding and potentially selects the outcome of Rho activation. Curr Biol 14, 1822–1826.

36. Parks, S, and Wieschaus, E (1991). The Drosophila gastrulation gene concertina encodes a G alpha-like protein. Cell 64, 447–458.

37. Perez-Mockus, G, Mazouni, K, Roca, V, Corradi, G, Conte, V, and Schweisguth, F (2017). Spatial regulation of contractility by Neuralized and Bearded during furrow invagination in Drosophila. Nat Commun 8, 1594.

38. Ratnaparkhi, A, and Zinn, K (2007). The secreted cell signal Folded Gastrulation regulates glial morphogenesis and axon guidance in Drosophila. Dev Biol 308, 158– 168.

39. Seher, TC, and Leptin, M (2000). Tribbles, a cell-cycle brake that coordinates proliferation and morphogenesis during Drosophila gastrulation. Curr Biol 10, 623–629.

40. Seher, TC, Narasimha, M, Vogelsang, E, and Leptin, M (2007). Analysis and reconstitution of the genetic cascade controlling early mesoderm morphogenesis in the Drosophila embryo. Mech Dev 124, 167–179.

41. Spahn, P, and Reuter, R (2013). A vertex model of Drosophila ventral furrow formation. PLoS One 8, e75051.

42. Sweeton, D, Parks, S, Costa, M, and Wieschaus, E (1991). Gastrulation in Drosophila: the formation of the ventral furrow and posterior midgut invaginations. Development 112, 775–789.

43. Urbansky, S, González Avalos, P, Wosch, M, and Lemke, S (2016). Folded gastrulation and T48 drive the evolution of coordinated mesoderm internalization in flies. Elife 5, e18318.

44. Vicente-Manzanares, M, Ma, X, Adelstein, RS, and Horwitz, AR (2009). Non-muscle myosin II takes centre stage in cell adhesion and migration. Nat Rev Mol Cell Biol 10, 778–790.

45. Xie, S, and Martin, AC (2015). Intracellular signalling and intercellular coupling coordinate heterogeneous contractile events to facilitate tissue folding. Nat Commun 6, 7161.

46. Xie, S, Mason, FM, and Martin, AC (2016). Loss of Gα12/13 exacerbates apical area dependence of actomyosin contractility. Mol Biol Cell 27, 3526–3536.

47. Yevick, HG, Miller, PW, Dunkel, J, and Martin, AC (2019). Structural Redundancy in Supracellular Actomyosin Networks Enables Robust Tissue Folding. Dev Cell 50, 586–598.e3.

48. Zusman, SB, and Wieschaus, EF (1985). Requirements for zygotic gene activity during gastrulation in Drosophila melanogaster. Dev Biol 111, 359–371.

